# Profiling DNA Methylation Differences Between Inbred Mouse Strains on the Illumina Human Infinium MethylationEPIC Microarray

**DOI:** 10.1101/194464

**Authors:** Hemant Gujar, Jane W. Liang, Nicholas C. Wong, Khyobeni Mozhui

**Affiliations:** Department of Preventive Medicine, University of Tennessee Health Science Center, Memphis, Tennessee 38163, USA; Monash Bioinformatics Platform, Monash University, Clayton VIC, Australia; Department of Genetics, Genomics and Informatics, University of Tennessee Health Science Centre, Memphis, Tennessee 38163, USA

**Keywords:** DNA methylation, epigenetics, microarray, cross-species comparison

## Abstract

The Illumina Infinium MethylationEPIC provides an efficient platform for profiling DNA methylation in humans at over 850,000 CpGs. Model organisms such as mice do not currently benefit from an equivalent array. Here we used this array to measure DNA methylation in mice. We defined probes targeting conserved regions and performed a comparison between the array-based assay and affinity-based DNA sequencing of methyl-CpGs (MBD-seq). Mouse samples consisted of 11 liver DNA from two strains, C57BL/6J (B6) and DBA/2J (D2), that varied widely in age. Linear regression was applied to detect differential methylation. In total, 13,665 probes (1.6% of total probes) aligned to conserved CpGs. Beta-values (β-value) for these probes showed a distribution similar to that in humans. Overall, there was high concordance in methylation signal between the EPIC array and MBD-seq (Pearson correlation r = 0.70, p-value < 0.0001). However, the EPIC probes had higher quantitative sensitivity at CpGs that are hypo-(β-value < 0.3) or hypermethylated (β-value > 0.7). In terms of differential methylation, no EPIC probe detected significant difference between age groups at a Benjamini-Hochberg threshold of 10%, and the MBD-seq performed better at detecting age-dependent change in methylation. However, the top most significant probe for age (cg13269407; uncorrected p-value = 1.8 × 10^−5^) is part of the clock CpGs used to estimate the human epigenetic age. For strain, 219 Infinium probes detected significant differential methylation (FDR cutoff 10%) with ∼80% CpGs associated with higher methylation in D2. This higher methylation profile in D2 compared to B6 was also replicated by the MBD-seq data. To summarize, we found only a small subset of EPIC probes that target conserved sites. However, for this small subset the array provides areliable assay of DNA methylation and can be effectively used to measure differential methylation in mice.

## Introduction

There has been a surge in large-scale epigenetic studies in recent years. In particular, epigenome-wide association studies (EWAS) of DNA methylation have shown association with physiological traits [1,2], diseases [3-5], environmental exposures [6,7], aging [8], and even socioeconomic [9] and emotional experiences [10]. The development of robust and reliable methylation microarrays has been an important driving force. In particular, the Illumina Human Methylation BeadChips have made it both convenient and cost-effective to incorporate an epigenetic arm to large epidemiological studies [11,12]. The latest version, the Illumina Infinium MethylationEPIC BeadChip (EPIC), provides an efficient high throughput platform to quantify methylation at 866,836 CpG sites on the human genome [13,14]. A remarkable biological insight that has emerged from these array-based studies is the definition of the methylation-based “epigenetic clock,” a biomarker of human age and aging (i.e., the epigenetic clock) that is defined using specific probes represented on these arrays [8].

Currently there is no equivalent microarray platform for model organisms and work in experimental species have largely relied on high-throughput sequencing. For instance, while the human DNA methylation age can be calculated from a few hundred probes on the Illumina BeadChips, a similar effort in mice required a more extensive sequencing of the mouse methylome [15]. However, CpG islands (CGIs) are largely conserved between mice and humans and the two species share similar numbers of CGIs and similar proportions of CGIs in promoter regions of genes [16]. Considering that these CpGs and CGIs are highly conserved in gene regulatory regions, it is feasible that probes on the human microarrays that target these sites may have some application in research using rodent models. This was previously evaluated for the two older versions of the Illumina HumanMethylation BeadChips [17]. The work by Wong et al. demonstrated that a subset of the probes targeting highly conserved sites provide reliable measures of DNA methylation in mice, and could be feasibly used to evaluate tissue specific methylation and in cancer related studies using the mouse as a model system.

In the present work, we extend the conservation analysis to the EPIC platform, and evaluate the capacity of these probes to detect differential methylation. We begin by defining the conserved probes and the key features of the corresponding CpG sites in the context of the larger mouse and human genome. We also compare the methylation signal detected by the conserved probes with affinity-based methyl-CpG enriched DNA sequence (MBD-seq) data from the same samples and evaluate if the conserved probes are informative of age and strain differences in mice.

## Materials and Methods

### Defining Conserved EPIC probes

Sequences for the 866,836 CpG probes were obtained from Illumina (http://www.illumina.com/). The probe sequences were aligned to the mouse genome (mm10) using bowtie2 (version 2.2.6) with standard default parameters. A total of 34,981 probes aligned to the mouse genome of varying alignment quality. Conserved probes were then defined based on quality of alignment. For this, we filtered out all sequences with a low mapping quality (MAPQ) of less than 60 (15,717 excluded) and those that contain more than two non-matching base pairs (1,092). To retain only the high quality probes, we further filtered probes based on confidence in DNA methylation signal and based on this, 4,507 probes with detection p-values > 0.0001 were removed. This generated a list of 13,665 high quality probes that are conserved sequences and provide reliable methylation assays in mice (these are listed in **Supplementary Data S1**). CpG island annotations [18] for the respective genome were downloaded from UCSC Genome Browser (http://genome.ucsc.edu) and distribution of conserved probes and positions of CGIs were plotted to the human (GRCh37) and mouse (mm10) genomes using CIRCOS [19].

For conserved sequences, there is high correspondence in functional and genomic features between mouse and human genomes and we referred to the human probe annotations provided by Illumina to define the location of conserved probes with respect to gene features and CpG context (i.e., islands, shores, shelves) (**Supplementary Data S1**). To evaluate if the conserved set is enriched in specific features relative to the full background set, we performed a hypergeometric test using the phyper function in R.

### Animals and sample preparation

Tissues samples were derived from mice that were part of an aging cohort maintained at the University of Tennessee Health Science Center (PI: Robert W. Williams). Details on animal rearing and sample collection are described in Mozhui and Pandey 2017 [20]. All animal procedures were approved by the UTHSC Animal Care and Use Committee.

Liver tissues were collected from mice aged at ∼4 months (mos; young), ∼12 mos (mid), and ∼24 mos (old). The mice were of two different strains—C57BL/6J (B6) and DBA/2J (D2)—and as the colony was set up to study aging in females, the majority of the mice in this study are females (**Table 1**). Mice were euthanized by intraperitoneal injection of Avertin (250 to 500 mg/kg of a 20 mg/ml solution), followed by cardiac puncture and exsanguination. All sample collection procedures were done on the same day within a 3-hour timeframe. Liver samples were snap-frozen and stored at -80°C until use.

**Table 1:** Sample details and average methylation signal intensity.

| Sample | Age | Age (months) | Strain<sup>1</sup> | Sex | Full set (850K) |  | Conserved set (13665) |  |
| --- | --- | --- | --- | --- | --- | --- | --- | --- |
|  |  |  |  |  | Mean | Median | Mean | Median |
| Mouse1 | young | 4 | D2 | F | 505 | 394 | 3206 | 1898 |
| Mouse2 | young | 4 | D2 | F | 926 | 524 | 10989 | 10278 |
| Mouse7 | young | 4 | B6 | F | 877 | 538 | 9866 | 8702 |
| Mouse8 | young | 4 | B6 | F | 766 | 397 | 10386 | 9975 |
| Mouse3 | mid | 12 | D2 | F | 852 | 483 | 10615 | 9880 |
| Mouse4 | mid | 12 | D2 | F | 818 | 430 | 10866 | 10542 |
| Mouse5 | mid | 12 | D2 | M | 845 | 456 | 11545 | 10982 |
| Mouse9 | mid | 12 | B6 | M | 852 | 444 | 11433 | 11187 |
| Mouse6 | old | 24 | D2 | F | 737 | 379 | 10206 | 9611 |
| Mouse10 | old | 24 | B6 | F | 845 | 448 | 10767 | 10436 |
| Mouse11 | old | 24 | B6 | F | 886 | 490 | 11302 | 10741 |
| Human1 |  |  |  |  | 7568 | 7218 | 8710 | 8616 |
| Human2 |  |  |  |  | 10668 | 10288 | 11761 | 11599 |

<sup>1</sup> D2: DBA/2J; B6: C57BL/6J

DNA was purified from the liver tissue using the Qiagen AllPrep kit (http://www.qiagen.com) on the QIAcube system. Nucleic acid quality was checked using a NanoDrop spectrophotometer (http://www.nanodrop.com). As reference, we also included two human samples. These are DNA derived from the buffy coats from two individuals.

### DNA methylation microarray and data processing

DNA methylation assays were performed as per the standard manufacturer’s protocol (http://www.illumina.com/). In brief, 500 ng of DNA extracted from the mouse liver was treated with sodium bisulfite to convert cytosine to uracil. The 5-methyl cytosine remains unreactive to sodium bisulfite. The DNA is then hybridized to the EPIC BeadChip. After washing off unhybridized DNA, a single base extension was recorded to calculate the methylation level at the CpG probe site. DNA methylation assays were performed at the Genomic Services Lab at the HudsonAlpha Institute for Biotechnology (http://hudsonalpha.org). Raw intensity data files (idat files) for both mouse and human samples were processed using the R package, Minfi [21].

The intensity and β-values were used to evaluate the performance of the EPIC probes in mice and humans. Comparisons were based on the full set of 850K probes and the conserved set of 13,665 probes. We also used the β-values and signal intensity scores for the 13,665 probes to perform hierarchical clustering and principal component analysis for the mouse samples. From initial quality checks, we identified one outlier mouse sample (**Supplementary Fig. S1**) that had lower intensity and higher detection p-value compared to the other mouse samples. This sample was excluded from the statistical tests.

### MBD-seq comparison

The mouse samples we report here were previously assayed for DNA methylation using MBD-seq [20]. This is an affinity-based enrichment of methylated CpGs using the methyl binding domain (MBD) of methyl-CpG-binding protein 2, followed by high throughput sequencing (MBD-seq) [22-24]. Sequencing was performed on Life Technologies’ Ion Proton platform. Data have been deposited to the NCBI’s Gene Expression Omnibus (https://www.ncbi.nlm.nih.gov/geo/; GEO accession ID GSE95361) and Sequence Repository Archive (https://www.ncbi.nlm.nih.gov/sra/; SRA accession ID SRP100703). To compare methylation signal detected by the conserved EPIC arrays, we extracted MBD-seq reads at the corresponding sites. MBD-seq does not provide single-base resolution as the resolution is limited to the fragment size, in this case ∼300 bp. However, since methylation levels at neighboring CpGs are largely correlated [25], we derived quantitative data from the number of read fragments that map to a CpG region. For the sites in the mouse genome targeted by the conserved EPIC probes, we expanded the window to 300 bp bins, and extracted the MBD-seq fragment counts. The CpG density-normalized methylation level was then quantified using the MEDIPS R package [26]. We then used Pearson’s correlation to compare the EPIC β-values and the relative methylation score (rms or the CpG density normalized methylation) detected by MBD-seq [27].

### Analysis of differential methylation

Statistical analyses were done in R (https://www.r-project.org/) and JMP Statistics (JMP Pro 12). Mice were grouped into three age categories (young, mid, and old; additional sample details are in **Table 1**). To evaluate differential methylation detected by the 13,655 conserved probes, we applied a regression model with age, strain and sex as predictors (∼ageGroups + strain + sex) for each probe using the R glm function and type III anova to calculate test statistics (equations are provided in **Supplementary Data S1**). For the MBD-seq reads, we performed differential methylation analysis of the read counts using the edgeR R package [28]. The same linear regression model was applied (∼ageGroups + strain + sex) and equations are provided in **Supplementary Data S1.** We then cross-compared differential methylation detected by the two methods. Treating the EPIC data as a discovery set, we applied the Benjamini-Hochberg (BH) procedure to control the false discovery rate (FDR) [29,30]. We then defined differentially methylated CpGs (DMCpGs) and evaluated the corresponding region in the MBD-seq data to test replication at a lenient uncorrected p-value threshold of 0.05. Likewise, in the reverse comparison, we applied an FDR threshold to identify differentially methylated regions (DMRs) in the MBD-seq data, and tested replication of the corresponding CpG at an uncorrected p-value threshold of 0.05.

## Results

### Conserved Infinium MethylationEPIC probes

The human EPIC array contains 866,836 50-mer probes. Out of these, we defined a total of 13,665 probes that align to conserved sites in the mouse genome and provide high quality methylation signal (details on mapping quality scores and methylation signal confidence are provided in **Supplementary data S1)**. In the full set of EPIC probes, 71% are located within annotated gene features or within 200–1,500 bp upstream of transcription start sites (TSS). Compared to this background set, a higher percent of the conserved probes (88%; 11,972 probes) target such functionally annotated regions. Probes that target CpGs located in exons, 5’ UTR, and within 200 bp upstream of TSS (TSS200) are highly overrepresented among the conserved set (**Table 2**). This is expected, since sequences in these functional regions are conserved across species. The upstream regulatory regions and the first exon harbor a large percent of CGIs, and compared to the background set, there is close to a 2.5-fold higher enrichment in CGIs among the conserved probes (**Table 2**). In contrast, there is no enrichment in probes that target CpGs that are between 200–1,500 bp upstream of TSS (TSS1500), gene body (mostly intronic), 3’ UTRs, and non-genic regions. Locations of the conserved probes and CGI densities in the human and mouse genomes are shown in **Fig. 1.**

**Table 2:** Genomic features of CpGs and enrichment in conserved sites.

|  | Full set<br>(850K) |  | Conserved set<br>(13665) |  |  |
| --- | --- | --- | --- | --- | --- |
| Feature | Counts | Percent<br>Total | Counts | Percent<br>Total | Enrichment p<sup>3</sup> |
| Gene features<sup>1</sup> |  |  |  |  |  |
| TSS1500 | 107193 | 12 | 1195 | 9 | ns |
| TSS200 | 65152 | 8 | 1940 | 14 | <1.0E-15 |
| 5'UTR | 73070 | 8 | 1269 | 9 | 1.8E-04 |
| 1stExon | 26433 | 3 | 2028 | 15 | <1.0E-15 |
| Exon | 5680 | 1 | 282 | 2 | <1.0E-15 |
| 3'UTR | 21594 | 2 | 340 | 2 | ns |
| Body | 318165 | 37 | 4918 | 36 | ns |
| Non-Genic | 249549 | 29 | 1693 | 12 | ns |
| CpG islands and flanking regions<sup>2</sup> |  |  |  |  |  |
| Islands | 161598 | 19 | 6270 | 46 | <1.0E-15 |
| Shores | 154735 | 18 | 2267 | 17 | ns |
| Shelves | 61811 | 7 | 664 | 5 | ns |
| Open Sea | 488692 | 56 | 4464 | 33 | ns |

<sup>1</sup> CpG position relative to gene features based on annotations from Illumina (UCSC\_RefGene\_Group). TSS1500 and TSS200 are CpGs at 0–200 or 200–1500 upstream of are transcription start sites; Non-genic are CpG with no annotated gene features.
<sup>2</sup> Shores = 0–2 kb from islands; shelves = 2–4 kb from islands
<sup>3</sup> Enrichment of gene features and CpG regions in the conserved set compared to the full set based on hypergeometric test

**Fig. 1.**
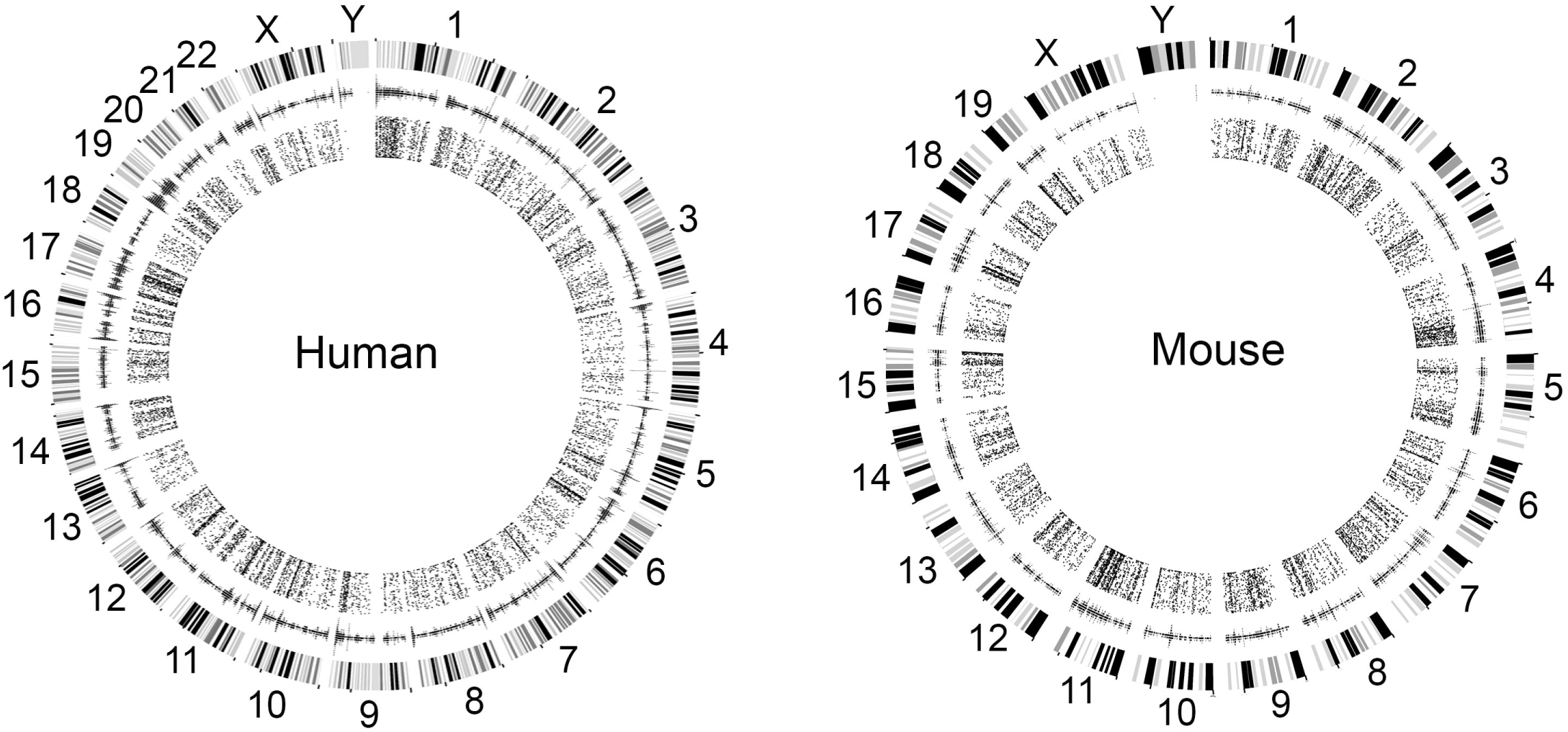
Location of conserved Illumina HumanMethylationEPIC probes and CpG densities in the human and mouse genomes. The outer circle displays the chromosomes and circular karyotype of the human and mouse genomes. CpG island (CGI) density is shown in the second circle. The innermost circle displays the positions of CpGs targeted by the 13,665 conserved probes.

### Comparison of probe performance in mouse and human samples

We used data generated from two human samples as reference. Using the full set of 850K probes, the mouse samples showed low overall signal intensity (**Fig. 2A**). The mean signal intensity for the two human samples was 9,118 ± 2,192 (**Table 1**). For the mouse samples, the mean signal intensity was 810 ± 114 (**Table 1**). The β-value distribution also showed poor performance for mice with a peak β-value at 0.4 that indicates failure for probes. The methylation β-values in human samples showed the expected bimodal distribution that characterizes the Illumina methylation arrays (**Fig. 2B**) [13,14].

**Fig. 2.**
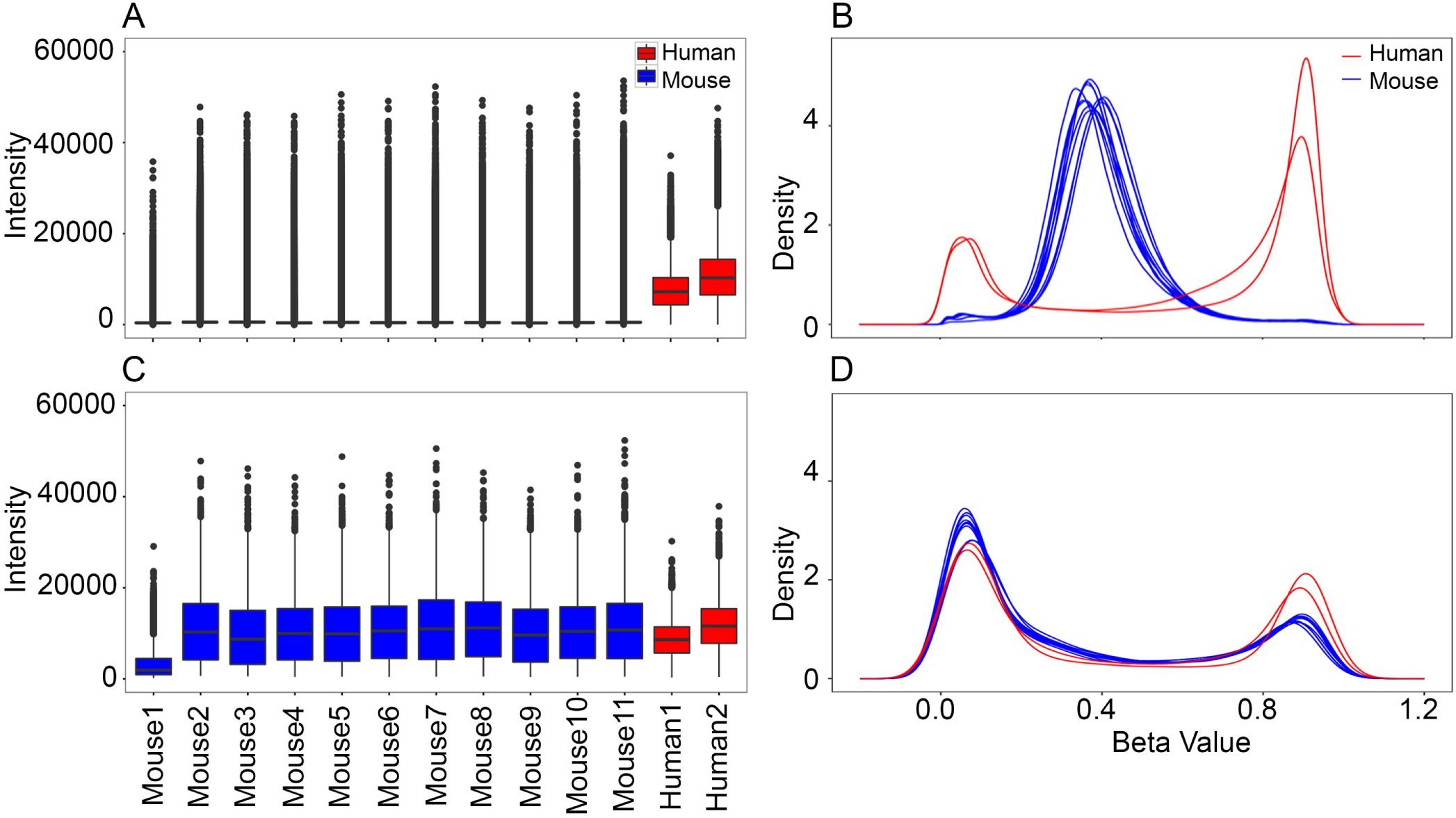
Distribution of signal intensities and methylation β-values in mice and humans. For the full set of 866,836 probes on the Illumina Infinium MethylationEPIC, the mouse samples have **(A)** low signal intensity compared to the two human samples, and **(B)** the β-values have a unimodal distribution that peaks at ∼0.4. The two human samples have the expected bimodal distribution for β-values. For the conserved set of 13665 probes, both the **(C)** signal intensity, and **(D)** the β-value distribution in the mouse samples are comparable to the two human reference samples. The signal intensity for mouse1 is relatively low for the conserved set of probes and this sample plots as an outlier in the principal component analysis.

The EPIC BeadChip clearly performed poorly in mice when we considered the full set of probes. However, when we considered only the 13,665 conserved probes, the methylation signal became comparable between the mouse and human samples. Total mean signal intensity for the mouse samples ranged from 9,866 to 11,545 (Mouse1, which failed the initial QC, has very low signal intensity compared to the other mouse samples; this was excluded from differential methylation analysis) (**Table 1**). Mean signal intensity for the two human samples were 8,711 and 11,761 (**Table 1**). The bimodal β distribution was also observed for this set of conserved probes in mouse samples (**Fig. 2C, 2D**).

### Comparison with MBD-seq

To determine if we could find concordant methylation signal, we compared the microarray β-values with the CpG density-normalized rms derived from MBD-seq data (average β-values and rms are provided in **Supplementary data S1**). Overall, there was concordance between the two technologies, and the β-values and rms were significantly correlated (Pearson’s correlation of 0.70, p < 0.0001). We grouped the EPIC probes into three categories based on β-values—hypomethylated for β < 0.3, hemimethylated for 0.3 ≤ β ≤ 0.7, and hypermethylated for β > 0.7—and examined correlations with the rms within each category. Given the high representation of islands and CpGs in 5’ regions of genes, which generally remain hypomethylated [16,31], the majority of the conserved probes fell into the hypomethylated category (**Table 3**). For the hypomethylated probes, 82% of the corresponding CpG regions also had rms < 0.3 (**Table 3**) and there was modest correlation between the rms and β-values (Pearson’s r = 0.18; p = 0.0001; **Fig. 3A**). For many of the CpGs regions that correspond to the hypomethylated probes, the rms were close to 0 and appeared unmethylated in the MBD-seq data. For hemimethylated probes, 58% of the corresponding regions had 0.3 ≤ rms ≤ 0.7 and 31% had rms < 0.3. This subset showed the highest correlation between the β-values and rms (r = 0.46; p < 0.0001; **Fig. 3B**). For hypermethylated probes, only 40% of corresponding regions were associated with rms > 0.7, and 54% had 0.3 ≤ rms ≤.7. This subset showed lower correlation between the β-values and rms (r = 0.04, p = 0.0392; **Fig. 3C**). The corresponding CpG regions for this hypermethylated set tended to have rms close to 0.75. This clustered rms distribution for CpG regions at the lower and upper levels of methylation indicate that the MBD-seq has lower quantitative sensitivity at these regions.

**Table 3.** Counts of Illumina HumanMethylationEPIC probes by β-value and concordance with MBD-seq at corresponding CpG regions.

| CpG Category<sup>1</sup> | Probes counts<sup>1</sup> | Counts of CpG regions by rms value<sup>2</sup> |  |  |
| --- | --- | --- | --- | --- |
|  |  | rms < 0.3 | 0.3 ≤ rms ≤ 0.7 | rms > 0.7 |
| Hypo<br>$\beta < 0.3$ | 7548 | 6198 | 1000 | 350 |
| Hemi<br>$0.3 \leq \beta \leq 0.7$ | 3159 | 973 | 1827 | 359 |
| Hyper<br>$\beta > 0.7$ | 2956 | 171 | 1599 | 1186 |

<sup>1</sup>Conserved probes on the HumanMethylationEPIC arrays were grouped by β-value. These are counts in each category.
<sup>2</sup>CpG For each category of probes, the corresponding CpG regions were counted and grouped by CpG density normalized relative methylation score (rms) to determine concordance between the array and MBD-seq

**Fig. 3.**
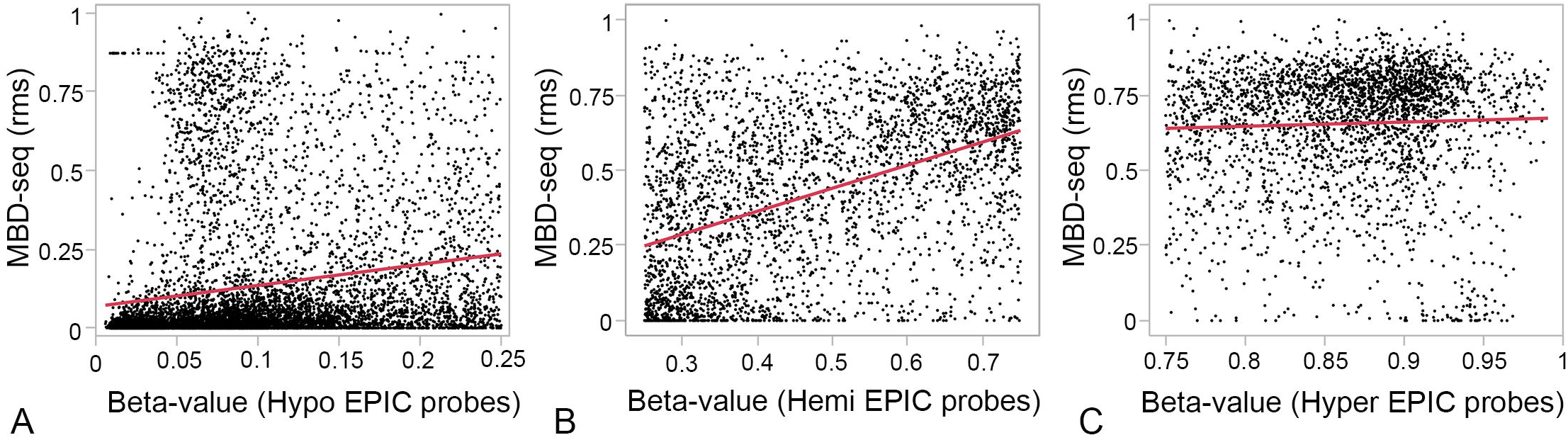
Correlation between MethylationEPIC and MBD-seq data. The 13,665 conserved MethylationEPIC probes were classified into three categories based on average β-values: hypo for β < 0.3, hemi for 0.3 ≤ β ≤ 0.7, and hyper for β > 0.7. For each probe, the 300 bp window around the corresponding CpG was determined and the CpG density-normalized relative methylation score (rms) was estimated for that region from MBD-seq data. A comparison between the β-values and rms showed **(A)** modest correlation for the hypomethylated CpGs (Pearson r = 0.18; p = 0.0001), **(B)** strong correlation for hemimethylated CpGs (r = 0.46; p < 0.0001), and **(C)** low correlation for hypermethylated CpG (r = 0.04, p = 0.0392). For CpGs with low β-values, the corresponding regions showed rms that cluster close to 0, and for CpG with high β-values, the corresponding rms tended to cluster close to 0.75.

Overall, the strong concordance with the MBD-seq data shows that the conserved EPIC probes provide a reliable quantification of methylation in mice. However, for CpGs that are hypomethylated or hypermethylated, the EPIC technology may have an advantage and provide higher quantitative sensitivity compared to the MBD-seq.

### Differential Methylation Analysis

We applied linear regression to examine differential methylation by age group and strain, and cross-referenced the DM-CpGs detected by the EPIC array with DMRs detected by MBD-seq. For the effect of age, no conserved EPIC probe passed a 10% FDR threshold (full results and p-values are provided in **Supplementary data S1**). However, we note that the probe that detected the most significant effect of age, cg13269407, is among the 353 CpGs that are used to estimate the human epigenetic age [8]. This CpG is hemimethylated (average β-value of 0.55) and associated with a ∼2.4-fold decline in methylation between young and old age (uncorrected p-value = 1.8 × 10^−5^). In the MBD-seq, the corresponding region is classified as hypomethylated with rms = 0 for most of the samples and no reliable statistics could be carried out for this region due to small number of mapped reads. We then performed a reverse comparison to identify age-dependent DMRs (age-DMRs) in the MBD-seq data and evaluated replication by the EPIC probes. At the same FDR threshold of 10%, the MBD-seq detected seven age-DMRs. These strong age-DMRs have rms between 0.3 and 0.7 and are associated with an increase in methylation with age. Most occur in CGIs that have been reported previously [20]. Out of these seven age-DMRs, six corresponding EPIC probes replicated the age-dependent increase in methylation at a nominal p-value cutoff of 0.05 (**Table 4**).

**Table 4.**
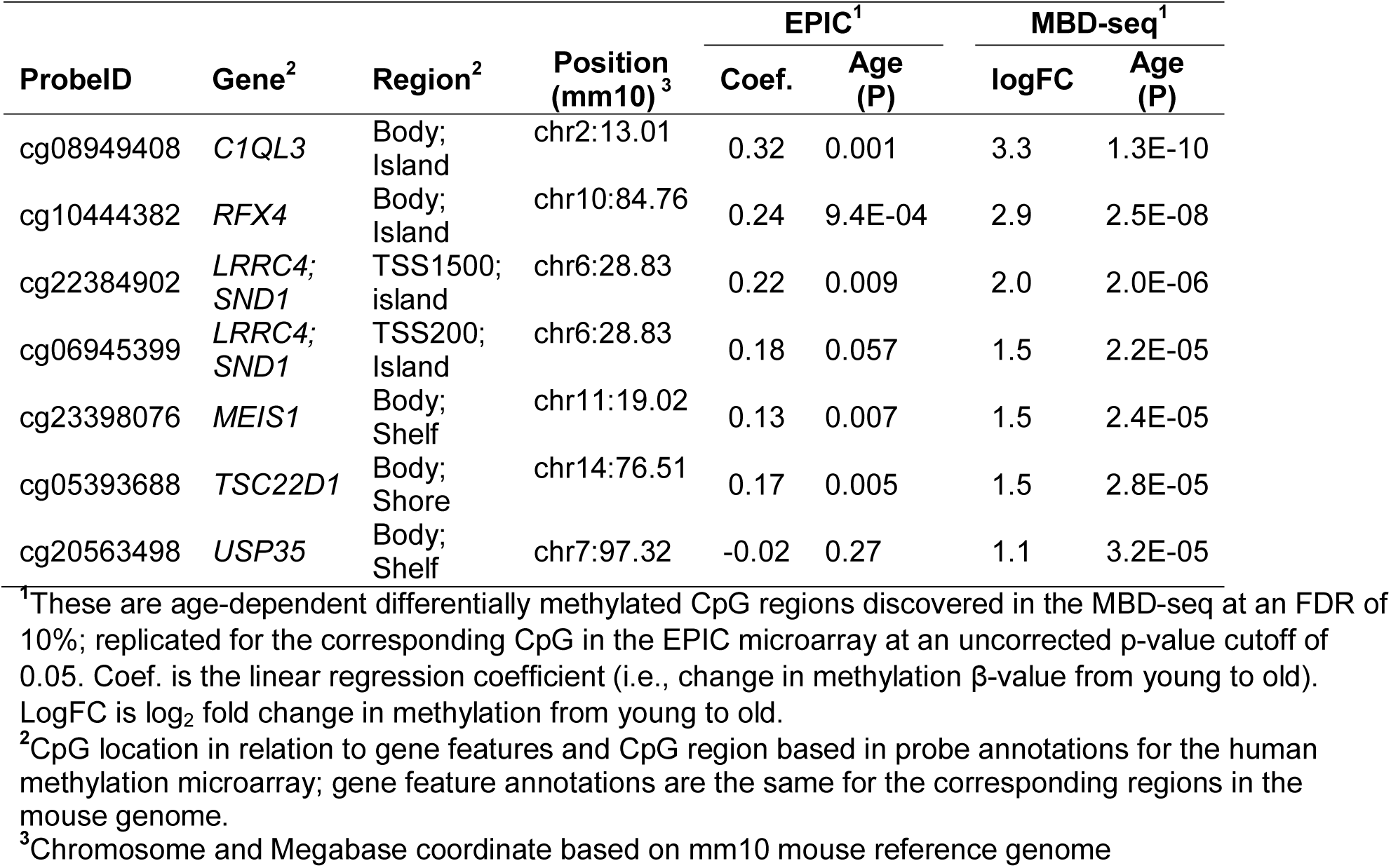
Age-dependent differentially methylated CpGs/regions detected by conserved probes and by MBD-seq.

For strain effect, 219 conserved EPIC probes detected significant difference in methylation between B6 and D2 at an FDR threshold of 10% (strain-DMCpGs). Close to 80% of these CpGs (175 out of 219) are associated with higher methylation in D2 relative to B6. In the MBD-seq data, only 29 of the 219 corresponding regions replicated strain effect at an uncorrected p-value cutoff 0.05 (**Table 5**). Of these, 9 were associated with higher methylation in B6, and 20 were associated with higher methylation in D2. In the reverse comparison, we identified only 37 strain-dependent DMRs (strain-DMRs) at an FDR cutoff of 10%. Consistent with the EPIC data, the majority of these regions (21 of the 37) showed higher methylation in D2 relative to B6. Of these, 16 strain differences were replicated at the corresponding CpG in the EPIC data (6 with higher methylation in B6 and 10 with higher methylation in D2) (**Table 5**).

**Table 5.**
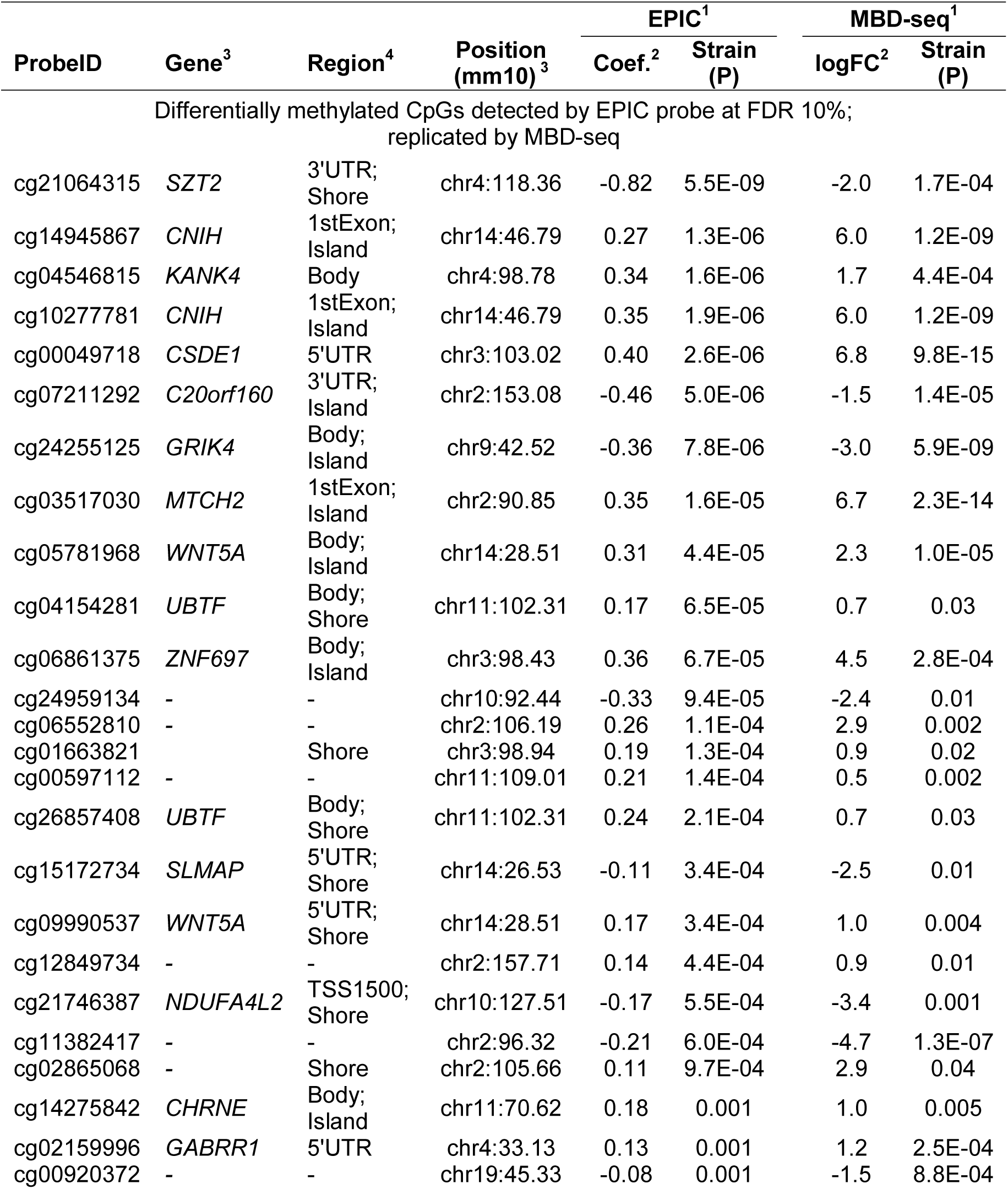
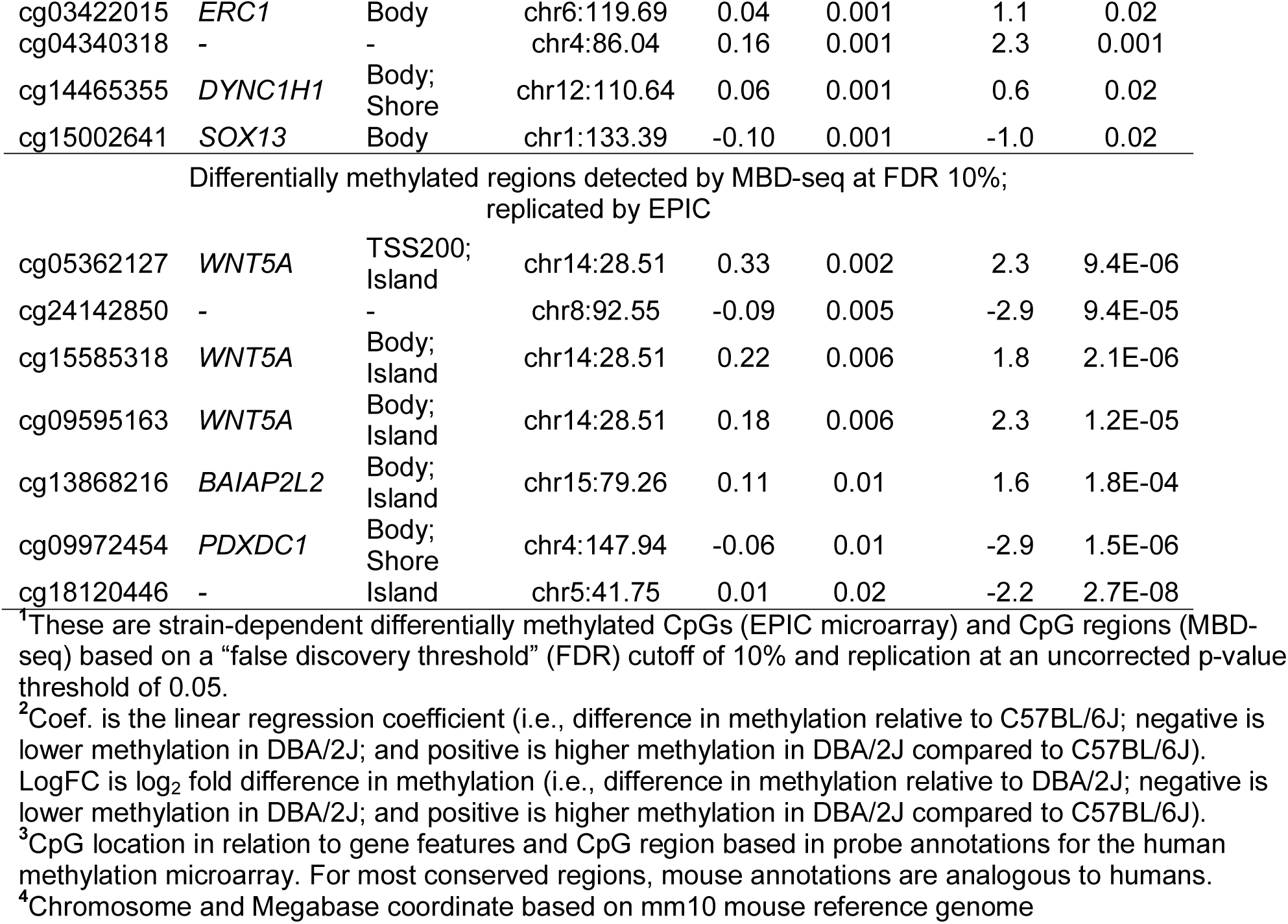
Strain-dependent differentially methylated CpGs/regions detected by both EPIC probes and by MBD-seq.

## Discussion

Given the high sequence conservation between the mouse and human genomes, we used the recently released Illumina MethylationEPIC microarray to assay DNA methylation at conserved CpGs in the mouse genome. We evaluated both the qualitative features as well as the quantitative performance and compared it with MBD-seq data that was generated on the same DNA samples from mice. Such a cross-species approach has been previously used to examine gene expression and perform comparative genomics studies [32-35]. The Illumina methylation array relies on bisulfite conversion and the probes are designed to target bisulfite-converted sequences. The two older versions of this Illumina methylation microarrays, the Infinium HumanMethylation 27K (HM27) and HumanMethylation 450K (HM450), have been carefully evaluated for use in mice [17]. The number of probes that map to the mouse genome can vary somewhat depending on the alignment algorithm. In the work by Wong et al. [17], alignment to the bisulfite-converted mouse genome resulted in the highest number of conserved probes. Using a stringent parameter of 100% sequence identity to the bisulfite genome, Wong et al. identified a total of 1,308 (4.7% of total) uniquely aligned probes in the 27K array, and 13,715 (2.8% of total) uniquely aligned probes in the 450K array that can be used to interrogate conserved CpGs in the mouse. In our present work, we performed alignment in a non-bisulfite space. While we required unique alignment, we tolerated up to two non-matching base pairs and added detection confidence as another parameter to identify probes that we can use for reliably quantitative assays. With these parameters, we identified 1.6% of total probes (13,665 in the 850K MethylationEPIC array) that aligned uniquely to the mouse genome and associated with high confidence in signal detection. In this set of 13,665 conserved EPIC probes, 9,429 (69%) were CpG loci carried over from the HM450 array and 7,483 of these were also in Wong’s list of conserved HM450 probes [17]. While alignment to the bisulfite-converted genome may have yielded a higher proportion of aligned probes, for our purposes the 13,665 probes provided a representative subset that we can use to assess quantitative performance in mouse samples and utility in detecting methylation variation.

The conserved probes mostly target CpGs located within annotated genes and in regulatory regions. In particular, exons, 5’ UTRs, CGIs in proximal regulatory regions (within 200 bp of TSS) are highly overrepresented among the set of 13,665 probes. This is expected since these coding and regulatory regions are the most conserved portions of the genome. Humans and mice have similar complements of CGIs and the genomic positions of these CGIs are also highly conserved, with 50% of CGIs located near annotated TSSs in both species [16,31,36]. In terms of quantitative variation in methylation, CGIs and promoter region CpGs show significant population variation [37]. However, compared to intergenic CpGs, the extent of inter-individual variability in methylation is reported to be much lower in these conserved sites [38,39]. Hence, an obvious limitation in using the conserved EPIC probes is that we attain only a narrow perspective of the mouse methylome and we may be sampling the portion of CpGs that shows the least quantitative variability in a population. Nonetheless, CpGs in regulatory regions and CGIs play crucial roles in development and cell differentiation, and are implicated in tumor development and aging [16,31,36,40,41]. While narrow in perspective, the conserved probes likely represent a subset of CpGs with high functional relevance and application in cross-species study of DNA methylation.

To evaluate the quantitative performance of the EPIC probes, we compared methylation levels measured by a complementary technology, MBD-seq. The type of methylation information measured by the microarray and MBD-seq are somewhat different. The EPIC probes, based on bisulfite conversion, measure the methylation status at a single CpG dinucleotide. MBD-seq, on the other hand, relies on affinity capture of DNA fragments by the methyl-CpG binding domain protein [22-24]. The affinity is directly proportional to the number of methylated CpGs in the DNA fragment and the methylation level is indirectly estimated based on the counts of sequenced reads that map to that region. This means that the resolution is inherently limited by the sizes of the fragments (in this case ∼300 bp). Since methylation of neighboring CpGs is generally correlated [42-44], MBD-seq provides information on the methylation level of CpGs in a region rather than one CpG site. For the 13,665 conserved EPIC probes, we extracted read counts from within 300 bp bins of the targeted CpGs and derived the CpG density-normalized read counts. Overall, there is strong concordance in methylation levels measured by the two technologies and the correlation between the β-values and rms was strongest for CpGs that are moderately methylated (we define these as β-values between 0.3 to 0.7 methylated). However, for CpGs that are hypomethylated and hypermethylated, the rms for the corresponding regions showed a more clustered distribution and indicated a limited quantitative sensitivity for MBD-seq and limited capacity in discerning quantitative variation at such CpG regions. Our observations agree with a previous study that compared HM450 and MBD-seq data generated using the same commercial kit we used [45].

For a direct comparison between the EPIC probes and MBD-seq, we applied the same regression model and crosschecked the DMCpGs and DMRs detected by the two technologies. While we expected a higher quantitative sensitivity for the EPIC probes as to age, the EPIC probes did not detect significant differential methylation at an FDR threshold of 10%. However, the topmost significant probe, cg13269407, is part of the 353 clock CpGs that are used to estimate human DNA methylation age [8]. Consistent with the negative correlation with age in humans, this age-informative CpG was associated with a ∼2.4-fold reduction in methylation in the old mice relative to the young mice. Aside from cg13269407, only 10 other human clock CpG probes were in the conserved set and none of these are associated with age in mice. Overall, the effect of age was weak when we considered individual CpGs. When we examined the corresponding CpG regions, the MBD-seq was more effective at detecting age-dependent methylation. At an FDR cutoff of 10%, we identified seven CpG regions that are classified as age-DMRs. These age-DMRs have been previously reported and show increases in methylation with age in mice [20]. For these age-DMRs identified by MBD-seq, we then checked whether the EPIC probes could verify the age effect. For this cross-verification, we used a less stringent statistical threshold of 0.05 for uncorrected p-values and found that six of the targeted CpGs are also associated with a significant age-dependent increases in β-values. Our observations suggest that age-dependent changes in methylation at these conserved sites may be more pronounced if we consider the correlated change of neighboring CpGs rather than methylation status of a single CpG. Despite the low overall quantitative sensitivity, the MBD-seq provides a complementary approach that may perform better for detecting methylation changes in regions harboring multiple correlated CpGs.

DNA methylation can vary substantially between mouse strains and a large fraction of this is likely due to underlying sequence differences between strains [20,46,47]. Strain variation in methylation has been shown to associate with complex phenotypes in mice such as insulin resistance, adiposity, and blood cell counts [48]. In our analysis, we detected 219 CpGs (i.e., 1.6% of the 13,365 interrogated CpGs) with a significant difference between strains at an FDR cutoff of 10%. A large majority (175 out of 219 CpGs) was associated with higher methylation in D2 compared to B6. While the overall lower methylation in B6 is intriguing, such variation between strains must be cautiously interpreted. It is well known that SNPs in probe sequences can have a strong confounding effect. This is particularly pernicious for mouse specific microarrays in which probe sequences are usually based on the B6 mouse reference, and as a result, there is more efficient hybridization for B6-derived samples, which results in a positive bias for this canonical mouse strain [49-51]. In the present work, since the EPIC array is based on the human sequence, we do not expect a systematic bias for one strain over the other. For replication, we referred to the MBD-seq data and only 29 out of the 219 corresponding CpG regions had consistent differential methylation between B6 and D2 in the MBD-seq.

Unlike using a human array that should not bias one mouse strain over another, the MBD-seq data is more vulnerable to technical artifacts caused by sequence differences. As is the general practice, we performed the alignment of the MBD-seq reads to the mouse reference genome. This means the alignment will be more efficient for sequences from B6, while sequences from D2 will have more mismatches. Since methylation quantification is estimated from the relative number of aligned reads, this may result in a systematic negative bias for D2, and methylation levels in regions with sequence differences will tend to have lower methylation due to poorer alignment. As a result, a higher fraction of strain-DMR will have lower methylation in D2 compared to B6 [20]. In the case that these conserved CpGs have higher methylation in D2 compared to B6, then the negative bias will lessen the quantitative difference between the strains. This may explain why the effect of strain is less pronounced in the MBD-seq data. In the MBD-seq, there were only 37 DMRs between B6 and D2 at an FDR threshold of 10%, and the EPIC probes replicated 16 of these. Out of the 37 strain-DMRs, the majority (21 of the 37) was associated with higher methylation in D2. Both the EPIC and MBD-seq therefore show an overall lower methylation profile in B6 compared to D2 that warrants further investigation and verification. Such strain differences in overall methylation has been previously reported for A/J and WSB/EiJ, with the A/J strain exhibiting higher methylation of CGIs in normal liver tissue compared to WSB/EiJ. This difference in the methylome was suggested to contribute to differential susceptibility for nonalcoholic fatty liver disease that characterizes the two strains [46]. In the case of B6 and D2, the two strains are highly divergent in a number of complex phenotypes ranging from behavioral and physiological to aging traits. The panel of recombinant inbred progeny derived from B6 and D2 (the BXD panel) has been used extensively in genetic research [52-56]. If there is indeed a distinct profile in DNA methylation between B6 and D2, then it will be of interest to evaluate if it segregates in the BXDs and how the methylome contributes to some of the phenotypic differences. The BXD panel could be an extremely rich and as yet untapped resource for methylome-wide analysis of complex traits that can then be integrated with the extensive systems genetics work that has already been done with this mouse family [57,58]. No doubt, large-scale analysis of genome-wide DNA methylation in mouse genetic reference panels will be greatly accelerated with the development of a mouse version of the Infinium methylation arrays. And as is the case with other types of arrays, it will be crucial that the probes are designed against a more diverse panel of strains so that investigators can derive a more unbiased readout of methylation [59].

To conclude, we have catalogued a small subset of EPIC probes that target conserved CpGs in the mouse genome and that provide reliable quantification of DNA methylation in mouse samples. While detection for age-dependent methylation was weaker for the EPIC probes compared to MBD-seq, we have identified significant strain variation in methylation at the conserved CpGs. Our results indicate lower methylation for B6 compared to D2 at sites that have significant strain effect. It is unclear how much of the strain variation results from underlying sequence differences between B6 and D2, and this strain-specific profile needs to be further evaluated and verified

## Acknowledgments

This work was supported by funds from UTHSC Faculty AwardUTCOM-2013KM. We are very grateful to Robert W. Williams for providing advice and access to the aging colony that is funded by NIA NIH grant R01AG043930.

